# Design of parallel *β*-sheet nanofibrils using Monte-Carlo search, coarse-grained simulations, and experimental testing

**DOI:** 10.1101/2023.11.09.565678

**Authors:** Sudeep Sarma, Tarunya Rao Sudarshan, Van Nguyen, Alicia S. Robang, Xingqing Xiao, Justin V. Le, Michael E. Helmicki, Anant K. Paravastu, Carol K. Hall

**Affiliations:** Department of Chemical and Biomolecular Engineering, North Carolina State University, Raleigh, NC 27695-7905, United States; Department of Chemical and Biomolecular Engineering, Georgia Institute of Technology, Atlanta, GA 30332, United States

## Abstract

Peptide self-assembly into amyloid fibrils provides numerous applications in drug delivery and biomedical engineering applications. We augment our previously-established computational screening technique along with experimental biophysical characterization to discover 7-mer peptides that self-assemble into “parallel β-sheets”, i.e., β-sheets with N-terminus-to-C-terminus *β*-strand vectors oriented in parallel. To accomplish the desired β-strand organization, we applied the *PepAD* amino acid sequence design software to the Class-1 cross-β spine defined by Sawaya et al. This molecular configuration includes two layers of parallel β-sheets stacked such that N-terminus-to-C-terminus vectors are oriented antiparallel for molecules on adjacent β-sheets. The first cohort of *PepAD* identified peptides were examined for their fibrillation behavior in DMD/PRIME20 simulations, and the top performing sequence was selected as a prototype for a subsequent round of sequence refinement. The two rounds of design resulted in a library of eight 7-mer peptides. In DMD/PRIME20 simulations, five of these peptides spontaneously formed fibril-like structures with a predominantly parallel *β*-sheet arrangement, two formed fibril-like structure with <50% in parallel *β*-sheet arrangement and one remained a random coil. Among the eight candidate peptides produced by PepAD and DMD/PRIME20, five were synthesized and purified. All five assembled into amyloid fibrils composed of parallel β-sheets based on Fourier Transform Infrared Spectroscopy, Circular Dichroism, Electron Microscopy, and Thioflavin-T fluorescence spectroscopy measurements.

## 1. INTRODUCTION

We seek to establish a workflow to design previously-unknown amino acid sequences to produce peptides that assemble into specific desired structures. It is known that peptides can self-assemble into architectures like nanofibers^1,2^, nanosheets^3^, nanotubes^4^, nanoparticles^5,6^, but understanding of the relationship between amino acid sequence and structures of assemblies is limited. Our ability to engineer supramolecular structures at the nanoscale impacts a wide variety of potential applications^7,8^ including as polymeric biomaterials^9^, tissue-engineering scaffolds^10–13^, hydrogels^2,8,14–17^, drug release agents^18,19^, and biomineralization components^20,21^. Short peptides are particularly desirable for biomaterials discovery because they can be easy to synthesize. Although peptide assemblies can be composed of molecules in various secondary structures, we focus here on β-sheet assemblies. “Bottom-up” strategies, in which the amino acid sequence length, composition and pattern are tailored, can be used to obtain supramolecular architectures with great structural variety and desired functional properties. The amino acid sequence composition of the peptides and their secondary structure drives the peptide self-assembly process to obtain peptide-based supramolecular assemblies.

In contrast to the “bottom up” search for amino acid sequences that we are aiming to establish here, previous designs for β-sheet peptide assemblies have been inspired by fragments from naturally-occurring amyloidogenic proteins, or have emphasized simple patterning of hydrophobic and polar amino acids. Examples of peptide fragments that self-assemble into β-sheet fibrils include the 7-mer peptide fragment Aβ (16-22) (sequence: *KLVFFAE*), which is associated with Alzheimer’s disease, and the fibril-forming segment of the yeast prion protein Sup35 (sequence: GNNQQNY). Furthermore, Lynn et. al. chemically modified Aβ (16-22) to assemble into nano-sheets^4,7^. Examples of peptides designed with hydrophobic/polar patterning include RADA16-I (sequence: Ac-RADARADARADARADA-NH_2_)^1,22^ and MAX1 (sequence: VKVKVKVKV^D^PPTKVKVKVKV-NH_2_)^2,23^, where Ac-indicates an acetylated N-terminus, -NH_2_ indicates an amidated C-terminus, ^D^P indicates proline with D-chirality, -V^D^PPT-corresponds to type-II’ β-turn to promote β-hairpin formation.

There are eight possible classes of 2-layer β-sheet structures, called cross-β structures spines, that peptides can form according to a 2017 paper by Sawaya and Eisenberg^24^. Although there have been significant advances in statistical biophysics and bioinformatics-based tools to predict amyloidogenic regions in a peptide sequence, “bottom-up” computational design of novel peptide sequences not known to adopt β-sheet-rich supramolecular structures is still a challenge.

In our previous work, we developed a workflow for computational and experimental discovery of 7-amino-acid peptides for self-assembly into amyloid structures. The chosen target structure was the Class-8 cross-β spine structure described by Sawaya et al^24^, with peptides arranged into a pair of stacked antiparallel β-sheets. The workflow started with PepAD; a Monte-Carlo based *pep*tide *a*ssembly *d*esign algorithm developed in the Hall lab. PepAD allows custom pre-settings for design parameters, such as the peptide length, amino acid sequence, backbone-scaffold, and hydration properties, to identify specific fibril-forming peptides. Additionally, PepAD uses atomistic force-fields rather than knowledge-based information and hence, enables the de novo design of peptides not known in nature. The self-assembling tendencies of the peptides identified by the PepAD algorithm are further evaluated using discontinuous molecular dynamics (DMD) simulations with the PRIME20 force field to examine their fibrilization kinetics. Eight of the twelve *in-silico* peptides identified by PepAD in our previous work successfully formed fibrils in the DMD simulations and self-assembled into anti-parallel β-sheets when tested experimentally^27^. Thioflavin-T (ThT) fluorescence measurements were used to monitor amyloid fibril formation. Peptide secondary structure was probed using circular dichroism (CD) spectroscopy and Fourier-transform infrared spectroscopy (FTIR). FTIR also reported on β-strand organization within β-sheets. Finally, fibrils were imaged using negative-stain transmission electron microscopy (TEM). All peptides tested in that study exhibited nanofiber formation with FTIR signatures of antiparallel β-sheets. Since the relative alignment of peptides in neighboring sheets was not examined via DMD simulation or experiment, we could not claim that these peptides should form Class 8 structures.

In this work, we sought to test the ability of the computational tools to design peptides with a different target β-strand organization, the Class-1 cross-β spine defined by Sawaya et. al ^24^. This structure contains a 2-layer β-sheet structure, with parallel-oriented β-strands in each layer and antiparallel-oriented β-strands between the two layers. Sawaya et. al^24^ reported 5 peptides that form this structure, including GNNQQNY of the prion protein Sup35. We know of no designer peptides and few naturally occurring peptides in this size range that assemble into parallel β-sheets. The energy associated with the hydrogen bond network of a parallel β-sheet is higher than that of an antiparallel β sheet^25^, which may be why antiparallel β-sheets are favored for peptides in this size range. In contrast, larger amyloid peptides tend to favor parallel β-sheets, which maximize overlap of hydrophobic residues on adjacent β-strands. Furthermore, one can use the simple heuristic that oppositely charged sidechains near opposite termini can favor antiparallel organization^26^, but we know of no analogous heuristic to favor parallel β-sheets. In the work described here, two rounds of designs were performed to obtain eight 7-mer peptides that self-assemble to form parallel β-sheets. In contrast to our workflow for antiparallel β-sheet structure, two rounds of PepAD design followed by DMD/PRIME20 simulations were needed to produce 8 candidate parallel β-sheet forming peptides for experimental testing. (As in our previous paper, the relative alignment of peptides in neighboring sheets was not analyzed.) Five of the peptides spontaneously formed fibril-like structures with a predominantly parallel *β*-sheet arrangement, two peptides formed fibril-like structures with <50% in parallel *β*-sheet arrangement and one peptide remained as a random coil in DMD/PRIME20 simulations. Fourier Transform Infrared Spectroscopy, Circular Dichroism, Electron Microscopy, and Thioflavin-T fluorescence spectroscopy measurements were conducted on 5 out of 8 peptides (commercially produced and received at >95% purity). These tests revealed that all 5 peptides self -assembled into parallel β-sheets.

## 2. RESULTS

### 2.1 First round of design of Class 1 cross *β*-spine forming peptides

PepAD is a Monte-Carlo based algorithm that searches for peptide sequences that can self-assemble to form supramolecular structures^27^. A score function, *Г_score_*, which considers (i) the binding free energy, Δ*G_binding_*, of the peptide chain with its neighboring peptides and (ii) the intrinsic self-aggregation propensity, *P_aggregation_*, of the individual peptides, is used to evaluate new peptide sequences. Details are provided in *Section 4.1*.

The PepAD algorithm requires an initial backbone scaffold to design peptide sequences that can self-assemble into the Class 1 cross-*β* spine. As mentioned earlier, Sawaya et. al reported in their study^24^ that the fibril forming segment GNNQQNY of the prion protein Sup35 forms a steric zipper characteristic of the Class 1 cross-*β* spine. Hence, GNNQQNY fibril was used as the reference peptide in the first round of design. This structure consists of a 2-layer amyloid fibril whose *β* strands are parallel within the *β*-sheet layer and antiparallel between the *β*-sheet layers (Figure 1a). Based on their study, we constructed two versions of the GNNQQNY structure that forms the Class 1 cross-*β* spine; these were used as starting backbone scaffolds in two parallel design rounds. Hereafter, we refer to these two initial backbone scaffolds as **Conf-1** and **Conf-2**.

**Figure 1.**
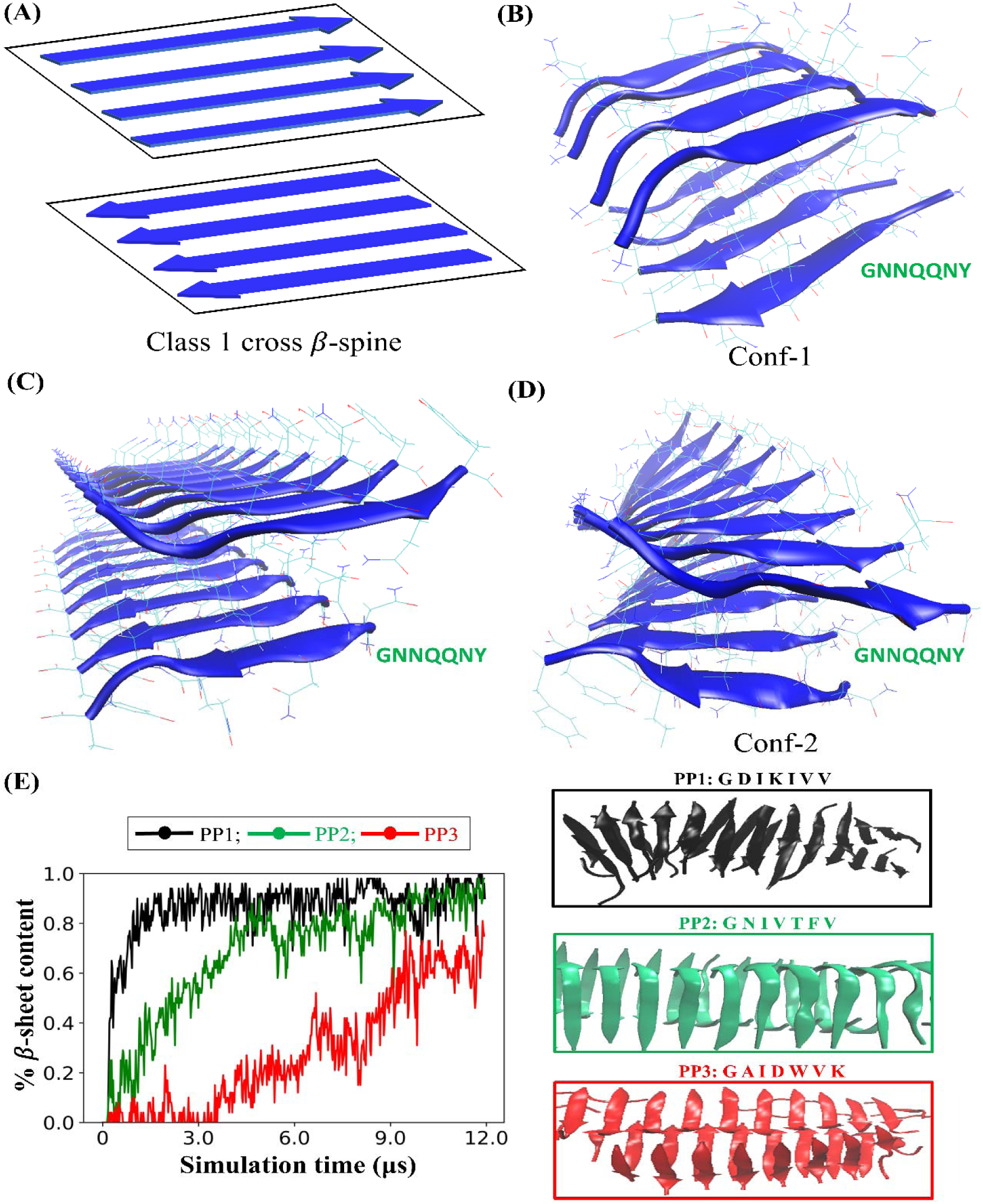
First round of design for Class 1 cross *β*-spine forming peptides: **(A)** Arrangement of peptides in the Class 1 cross *β*-spine, a 2-layer amyloid fibril consisting of parallel-oriented β-strands in each layer and antiparallel-oriented β-strands between the two layers. **(B)** Input fibril structure of peptide GNNQQNY for the PepAD algorithm (**Conf-1**) constructed using Discovery Studio 3.5 and Packmol packages and optimized by atomistic MD simulation in the AMBER14 package. **(C)** Fibril structure of peptide GNNQQNY obtained from PDB structure: 2omm (**D)** Input fibril structure of peptide GNNQQNY for the PepAD algorithm (**Conf-2**) constructed by performing a 25 ns MD simulation on structure in (C). **(E)** Plot of *β*-sheet content v/s simulation time describes the self-aggregation kinetics of peptide PP1 (sequence: GDIKIVV), PP2 (sequence: GNIVTFV) and PP3 (sequence: GAIDWVK) (left). The snapshot of the final simulation structures of peptides PP1, PP2, and PP3 indicating its fibrilization behavior is shown (right).

To obtain **Conf-1**, we constructed a 2-layer amyloid *β*-sheet structure using the Discovery Studio 3.5 and Packmol packages. Eight 7-mer GNNQQNY (PP0) peptide sequences were aligned in a parallel arrangement within each *β* sheet layer and an antiparallel arrangement between the *β* sheet layers with 4 peptides in each layer. The peptide distance within each sheet was set to be ∼ 5.5 Å and the inter-sheet distance was specified as ∼ 12 Å. A 5 ns explicit-solvent atomistic molecular dynamics simulation was conducted using the AMBER14 package to relax the two-layer *β*-sheet structure in the afore-mentioned parallel arrangement and to eliminate any atomic overlaps. The structure obtained following the 5ns simulation was used as the input backbone scaffold (**Conf-1**) for the PepAD algorithm (Figure 1b).

To build **Conf-2**, we used the crystal structure of peptide GNNQQNY (PDB ID: 2omm) reported in the Protein Data Bank. The primary coordinate file corresponding to PDB ID: 2omm contains the crystal asymmetric unit of a single GNNQQNY sequence and provides the information needed to generate the biological assembly of four GNNQQNY peptides into a Class 1 cross-*β* spine. This is a 2-layer *β*-sheet structure containing 2 parallel-oriented *β*-strands in each layer with antiparallel oriented *β*-strands between the two layers. An in-house python code was written to generate a configuration of the Class 1 cross-*β* spine with 4 peptides (Supplementary Figure S1). Four replicas of this structure were generated to produce a 2-layer *β*-sheet structure with 8 parallel-oriented *β*-strands in each layer and an antiparallel arrangement between the two layers (Figure 1c). (Code is available at: https://github.com/CarolHall-NCSU-CBE/Parallel-self-assembling-peptides). We performed a 5 ns explicit solvent atomistic molecular dynamics simulation using the AMBER14 package to relax the aforementioned 2-layer sheet structure to eliminate any atomic overlaps. The structure obtained following the 5ns simulation was used as a second input backbone scaffold (**Conf-2**) for the PepAD algorithm (Figure 1d).

Next, we specified the hydration properties for the designed amyloid-forming peptides as input parameters for the PepAD algorithm. Two cases were investigated with two different sets of hydration properties for the peptide chain. We classify the 20 natural amino acids into four residue types: hydrophobic residues (Leu, Val, Ile, Ala, Met, Phe, Tyr, Trp, Gly), polar residues (Ser, Thr, His, Asn, Gln), charged residues (Arg, Lys, Asp, Glu) and other residues (Cys, Pro). The two cases are as follows, Case 1: *N*_hydrophobic_ = 5, *N*_polar_ = 0, *N*_charge_ = 2, *N*_other_ = 0 and Case 2: *N*_hydrophobic_ = 5, *N*_polar_ = 2, *N*_charge_ = 0, *N*_other_ = 0. (We were interested in determining which hydration properties favor the parallel amyloid-*β* sheet formation.) For each case we performed the PepAD algorithm at two different values for the weighting factor λ, viz. λ=2.0 and λ=3.0 in equation (1). The weighting factors λ=2.0 and λ=3.0 were chosen to provide a good balance between optimizing the binding free energy (Δ*G_binding_*), and the aggregation propensity terms (*P_aggregation_*) in the *Г_score_* of the amyloid-forming structure. All the searches start with random peptide sequences draped on the fixed backbone scaffold. By having random initial peptide sequences, we encourage our designs to proceed along different search pathways in sequence space and thereby sample peptides from a larger pool of peptide sequences than would otherwise be the case. The ***Г**_score_* profile fluctuates considerably as new amino acids are placed on the different sites of the peptide chain. By examining the ***Г**_score_* profiles over the sequence evolution, we identified four peptide sequences (PP1-PP4) with low scores for evaluation using DMD/PRIME20 simulations. PP1 and PP2 were identified with **Conf-1** as the starting structure, and PP3 and PP4 were identified with **Conf-2** as the starting structure.

We performed a preliminary screen to investigate the fibrilization kinetics of the four PepAD identified *in-silico* peptides (PP1-PP4) using DMD/PRIME20 simulations. DMD/PRIME20 is a fast alternative to traditional molecular dynamics simulations that uses discontinuous potentials to model peptide aggregation. The force field PRIME20, developed by the Hall group in 2010, is a coarse-grained model where each amino acid is represented by a 3-sphere backbone comprised of united atoms (NH, CαH and CO) and a single-sphere side chain, R^28–33^. Details are provided in *Section 4.2*. The simulations were performed at temperatures ranging between 296 K to 310 K for 5 μs. The peptides were then extensively studied at the temperature at which they showed the highest fibrillation propensities by performing 12 μs DMD/PRIME20 simulations. The simulations predict that peptide PP1 (sequence: GDIKIVV), PP2 (sequence: GNIVTFV) and PP3 (sequence: GAIDWVK) spontaneously form amyloid-like fibrils and adopt a predominantly parallel *β*-sheet arrangement. Peptide PP4 (sequence: GGIDWKI) formed amyloid-like fibrils but exhibited less than 50% parallel *β*-sheet content in the DMD/PRIME20 simulations. Table 1 contains the sequences of peptides PP1-PP4 with their associated scores, binding free energies, intrinsic self-aggregation propensities; the number of layers in the fibrils and the parallel *β*-sheet content percentage were estimated from DMD/PRIME20 simulations. The % *β*-sheet content v/s simulation time for peptides PP1-PP3 and PP4 are shown in Figure 1E (left) and Supplementary Figure S2, respectively. (Snapshots of the final simulated structures of peptides PP1-PP3 are shown in Figure 1E (right).

**Table 1.** The sequences of the four *in-silico* discovered peptides in the first round of design with their corresponding *Г_score_*, Δ*G_binding_*and λ *⨉ P_aggregation_* values computed from PepAD. The %Parallel *β*-sheet content for each peptide is computed from the DMD/PRIME20 simulations.

| Peptides | Case | Sequences | $\Gamma_{score}$ | $\Delta G_{binding}$ | $\lambda \times P_{aggregation}$ | DMD/PRIME20 |
| --- | --- | --- | --- | --- | --- | --- |
| Starting with Conf-1 |  |  |  |  |  |  |
| PP1 | 1 | GDIKIVV | -12.19 | -12.37 | -0.18 | Two-layer fibril, ~77% parallel β-sheet content |
| PP2 | 2 | GNIVTFV | -17.25 | -12.39 | 4.85 | Two-layer fibril, ~100% parallel β-sheet content |
| Starting with Conf-2 |  |  |  |  |  |  |
| PP3 | 1 | GAIDWVK | -8.28 | -16.20 | -7.92 | Multi-layer fibril, ~100% parallel β-sheet content |
| PP4 | 1 | GGIDWKI | -6.55 | -15.88 | -9.33 | Two-layer fibril, ~46% parallel β-sheet content |

### 2.2 Second round of design for Class 1 cross *β*-spine forming peptides

Since peptide PP2 (sequence: GNIVTFV) was the most promising candidate in our first round of design of Class 1 cross-*β* spine forming peptides when studied via DMD simulations, it was selected as the reference sequence to create the initial backbone scaffold to perform a second round of *in-silico* peptide design. (We liked that the *β*–sheets which assembled in the DMD simulations were 100% parallel.) To build the backbone scaffold we used the Pymol software to mutate the residues on the Class 1 cross-*β* spine structure of 16 GNNQQNY peptides (Figure 1C) to generate a Class 1 cross-*β* spine structure containing 16 GNIVTFV (PP2) peptides (Supplementary Figure S3). For best comparison with the DMD/PRIME20 simulations and experimental biophysical characterization (*see Section 4.3*), the N-termini and C-termini were acetylated and amidated (N-cap and C-cap), respectively in this round of design using the PepAD algorithm. (The main effect of “patching” of termini is to eliminate charges at neutral pH). A 25ns explicit-solvent atomistic molecular dynamics simulation was conducted using the AMBER14 package to relax the parallel 2-layer PP2 *β*-sheet structure and eliminate any atomic overlaps. The structure obtained following the 25ns simulation (Figure 2(A)) was used as the input backbone scaffold for the second round of PepAD design. We refer to this structure as **Conf-3**.

**Figure 2.**
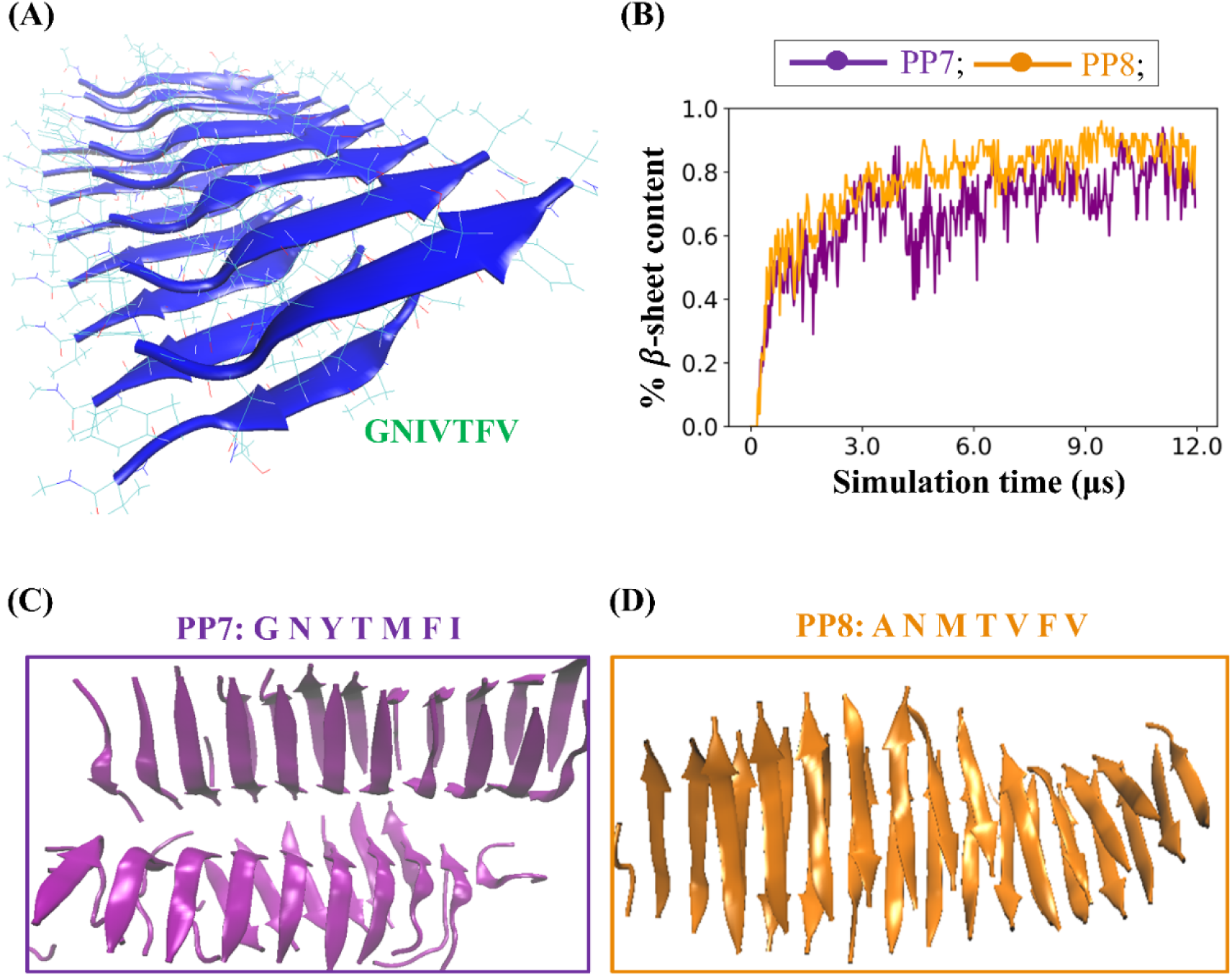
Second round of design of Class 1 cross *β*-spine forming peptides: **(A)** Input fibril structure of peptide PP2: GNIVTFV (with patched N- and C-termini) for the PepAD algorithm in Round 2 of design (**Conf-3**) (**B)** Plots of *β*-sheet content v/s simulation time describe the self-assembly kinetics of peptides PP7 and PP8. Snapshots of the final structures of **(C)** PP7 and **(D)** PP8 were obtained from the DMD simulations.

We next performed the PepAD algorithm to generate a new cohort of parallel amyloid-forming peptides with Case 1 and Case 2 hydration properties using peptide PP2 draped on **Conf-3** as the reference peptide. Four peptides (PP5-PP8) obtained from this round were further investigated in DMD simulations to study their fibrillation kinetics. We again performed DMD simulations of the peptide systems for 5 μs to examine their self-aggregation kinetics at temperatures ranging from 296.1 K to 310K. These peptides were then extensively studied at the temperatures at which they showed the highest fibrillation propensity for 12 μs in DMD/PRIME20 simulations. Our simulations revealed that peptides PP5 (sequence: ADKVMFV) and PP7 (sequence: GNYTMFI) exhibited high parallel *β*-sheet content while peptide PP8 (sequence: ANMTVFV) exhibited < 50% parallel *β*-sheet content. Peptide PP6 (sequence: GDFVKFV) predominantly remained as random coil in our DMD/PRIME20 simulations. The sequence of peptides PP5-PP8 with their associated scores, binding free energies and intrinsic self-aggregation propensities, and observations from DMD/PRIME20 simulations are reported in Table 2. The % *β*-sheet content v/s simulation time of PP7 and PP8 obtained from DMD/PRIME20 simulations are shown in Figure 2B. Snapshots of the final simulated structures of peptides PP7 and PP8 are shown in Figures 2C and 2D respectively.

**Table 2.**
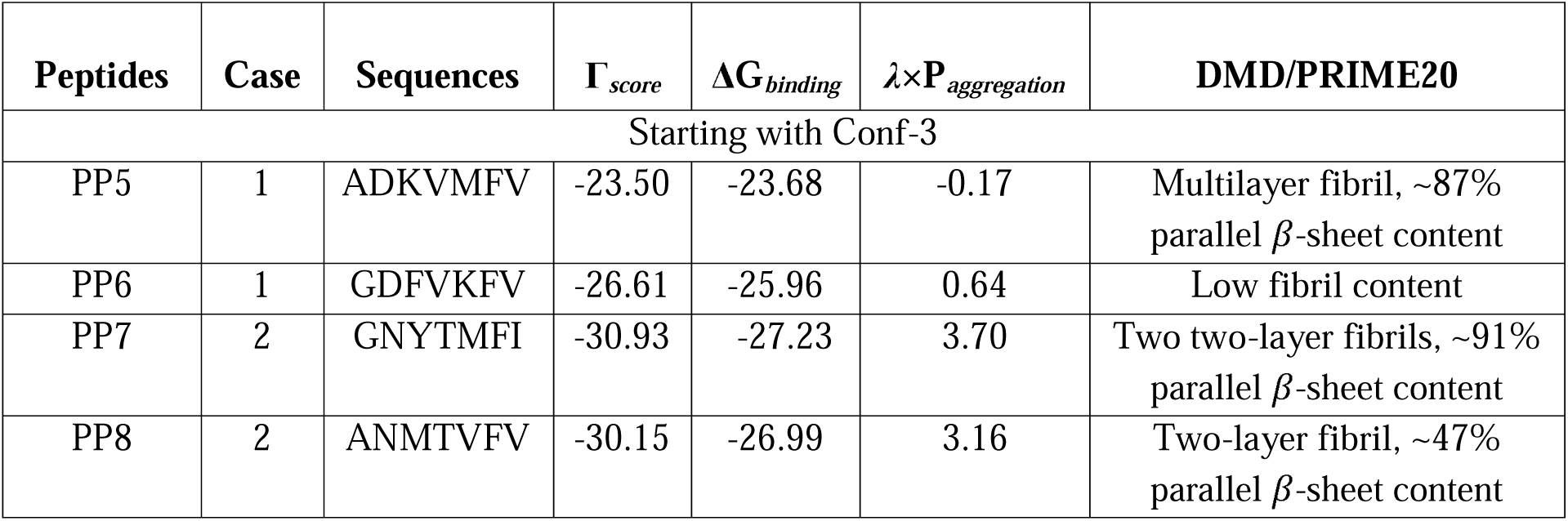
The sequences of the four *in-silico* discovered peptides obtained from the second round of design with their corresponding *Г_score_*, Δ*G_binding_* and λ *⨉ P_aggregation_*values computed from PepAD. The %Parallel *β*-sheet content for each peptide is computed from the DMD/PRIME20 simulations.

### 2.3 Experiments revealed parallel beta-sheets nanofibers

We experimentally evaluated the assembly of 5 out of the 8 candidates in Tables 1 and 2. We ordered commercial production of PP1, PP2, PP3, PP4, PP7 and PP8 (note that PP8 exhibited low fibrillar content in the DMD/PRIME20 simulations), but PP2 was not experimentally tested as it did not meet purity standards of 95%. Peptides PP5 and PP6 were not synthesized. All five final sequences formed parallel *β*-sheet nanofibers as evidenced by biophysical characterization techniques like negative-stain Transmission Electron Microscopy (TEM), Fourier-transform infrared spectroscopy (FTIR), Thioflavin-T Fluorescence (ThT) and Circular Dichroism (CD). All 5 sequences exhibited parallel *β*-sheet assembly.

The experimental techniques that we employed tend to be compatible with varying peptide concentrations. We performed Fourier-transform infrared spectroscopy at a concentration of 10mM, Transmission Electron Microscopy at a concentration of 1mM, Circular Dichroism at a concentration of 0.2mM, and Thioflavin-T Fluorescence at a concentration of 1mM. It is important to note that concentration of peptide solutions is a factor in the formation of *β*-strands. This concentration difference may result in varying levels of assembly between experimental techniques.

We observed the most definitive evidence of assembly and parallel *β*-sheets using FTIR and TEM, which we performed on peptide solutions at the highest peptide concentration (10mM). TEM can image the presence of nanofiber peptide assemblies. FTIR is sensitive to β-sheet secondary structure and can differentiate between parallel and antiparallel β-sheet structures. An FTIR peak near 1620cm^−1^ indicates β-strand secondary structures and an additional peak near 1690cm^−1^ is attributed to antiparallel organization of β-strands within the β-sheet. Figure 3A is a TEM image of fibrils of peptide PP3 showing assembly. Supplementary Figure S4 shows TEM images of all peptides. TEM Images reveal that all peptides form fibrils with thicknesses consistent with multi-layer *β*-sheets. Figure 3B compares the FTIR spectra from peptide PP3 to peptide P12 (sequence: ALRLELA) from our previous work^27^. P12 is a peptide from our previous successful effort to design peptides to form antiparallel β-sheets. The spectra of both peptides show a peak intensity near 1620cm^−1^ (1619cm^−1^ and 1625cm^−1^ respectively) indicating *β*-strand secondary structure, as expected. The additional peak at 1690cm-1 observed for P12 but not PP3 is indicative of anti-parallel organization of *β*-sheets. Figure 4 compares FTIR spectra from all peptides tested in our present work (Figure 4a) to all peptides tested in our previous work (Figure 4B)^27^. The former group of peptides did not exhibit a peak at 1690cm^−1^, whereas the latter group of peptides did. We interpret these results to indicate that the established design and characterization workflow was successful in producing self-assembling peptides with the target parallel *β*-strand organization.

**Figure 3.**
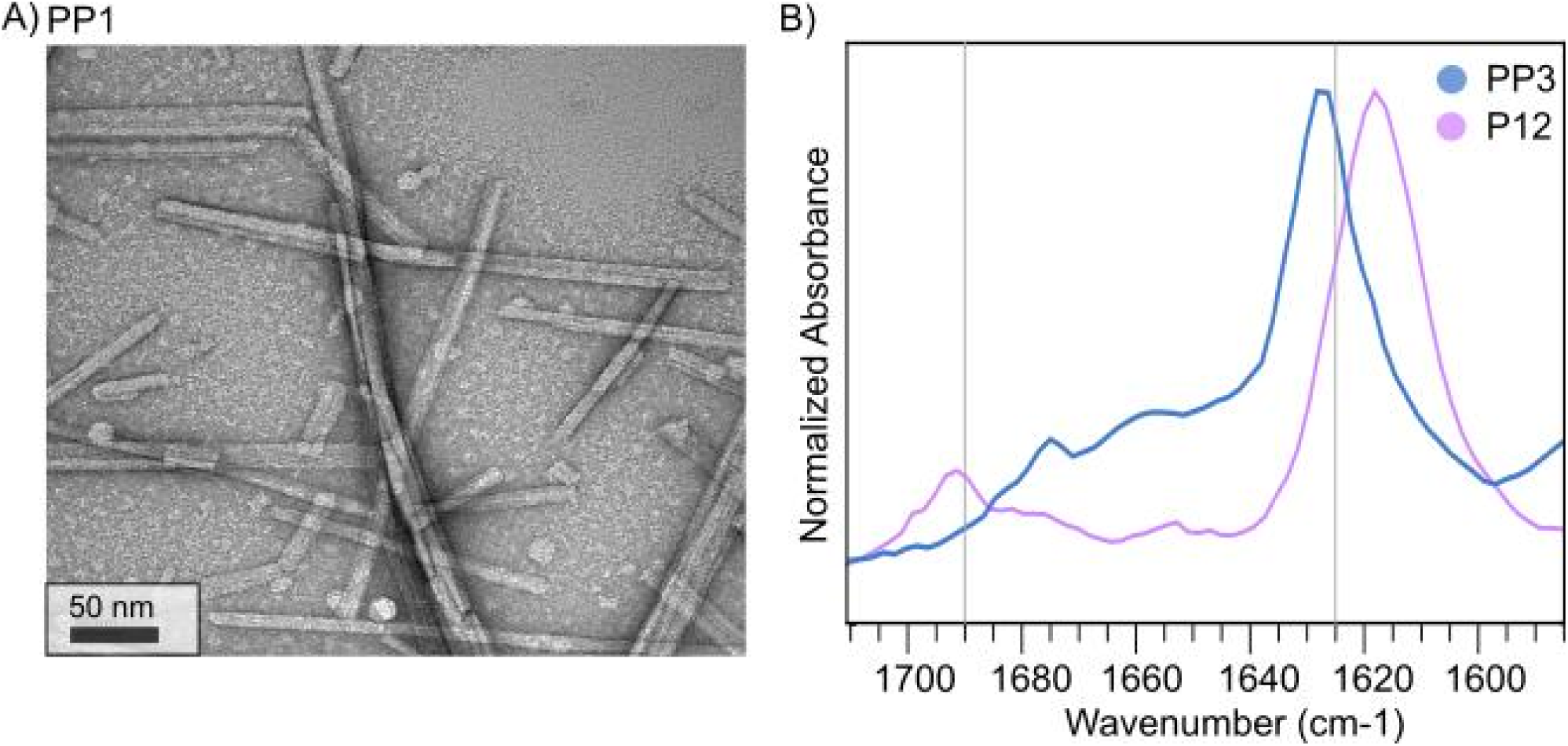
TEM Imaging and FTIR spectra **(A)** TEM Image of PP1 (**B)** FTIR spectra comparing P12 from our previous work and PP3. Note that PP3 shows an absence of a peak at 1690cm^−1^, which corresponds to antiparallel β-sheets.

**Figure 4.**
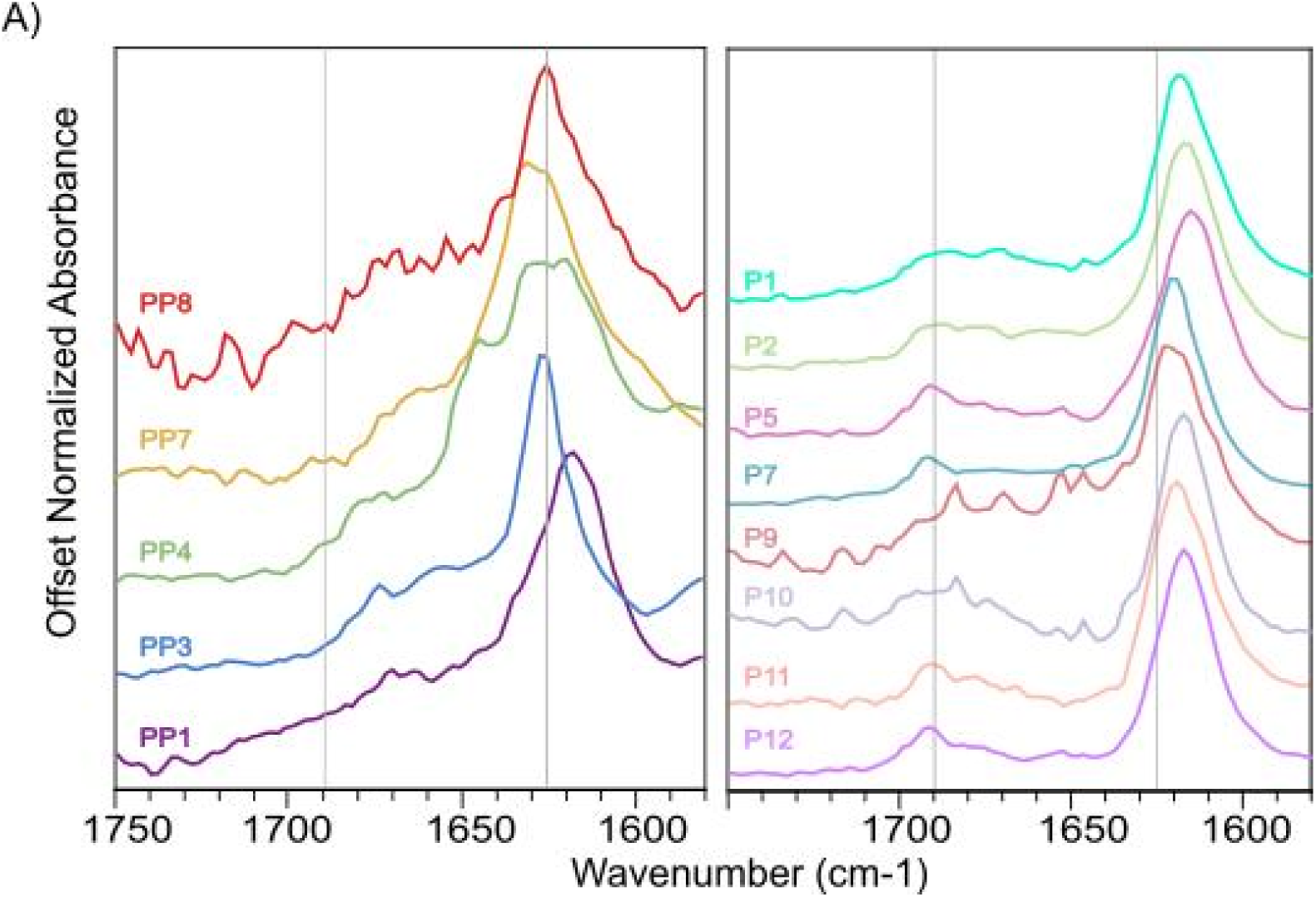
FTIR Spectra comparing peptides in our present study (left) with peptides from our previous study (right). Note that the peak at 1690 cm^−1^, associated with antiparallel β-sheets, is absent in spectra on the left.

We used Thioflavin-T fluorescence measurements to probe kinetics of β-sheet formation at peptide concentrations of 1mM. Figure 5A indicates that PP7 and PP8 assemble immediately, while PP3 and PP4 show low but increasing levels of fluorescence over 72h. PP1 has low fluorescence levels but shows evidence of assembly through other tested experimental methods. Note that absolute fluorescence level cannot be readily compared between different peptides: they are affected by factors such as binding of thioflavin-T to β-sheet surfaces and conformation of bound ThT molecules, which both depend on amino acid sequence. There are differences in the kinetics of assembly among these peptides. PP3 and PP4 both show a gradual increase in fluorescence during Thioflavin-T fluorescence experiments indicating slower assembly than PP7 and PP8. In fact, definitive FTIR spectra for PP3 and PP4 were obtained only after 6 days post assembly. In contrast PP7 and PP8 assemble immediately during Thioflavin-T Fluorescence experiments. This variation in kinetics was not observed in our previous study of peptides that form antiparallel *β*-sheets.

**Figure 5.**
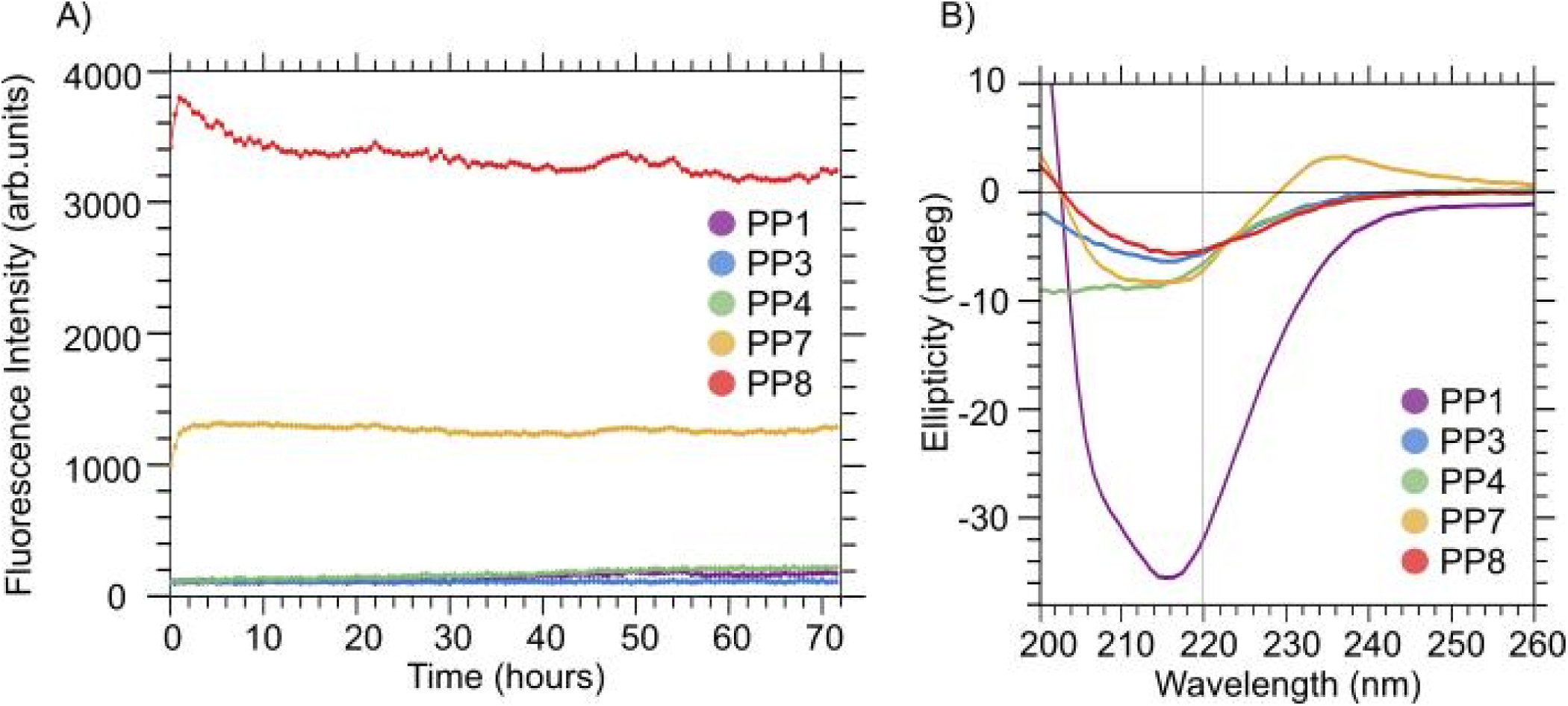
Thioflavin-T Fluorescence and Circular Dichroism **(A)** Fluorescence intensity plotted with time for peptides in our present study **(B)** Circular Dichroism experiments indicate that there is a dip at ∼215nm characteristic of β-sheet secondary structure.

We performed Circular Dichroism (CD) experiments on the peptides to characterize their secondary structure at 0.2mM. Assembled *β*-sheets are expected to show a single minimum near 220nm in CD spectra. We observed CD spectra consistent with heterogeneous secondary structure with partial *β*-strand signatures (Figure 5B). We used the BeStSel (Beta Sheet Selection) web server-based tool to predict the secondary structural distribution based on the CD spectra (Supplementary Table S1)^34^. BeStSel interpreted the spectrum of PP1 to correspond to 100% β-sheets, whereas it associated the other CD curves to between 30 and 65% β-strands. BeStSel also attempts to distinguish between parallel and antiparallel β-sheets. In agreement with our FTIR results in Figure 4, BeStSel attributed parallel β-sheet structure of PP1. However, for peptides PP3, PP4, PP7, and PP8, BeStSel assigned antiparallel β-sheet structure to the partial β-strand signatures, which is inconsistent with our FTIR results. We suggest that BeStSel is unreliable in its ability to distinguish between parallel and antiparallel β-strand organization, especially when secondary structure is heterogeneous. Table 3 summarizes the results of DMD/PRIME20 simulations and experimental measurements (FTIR, ThT, TEM and CD) of peptides PP1-PP8.

**Table 3.**
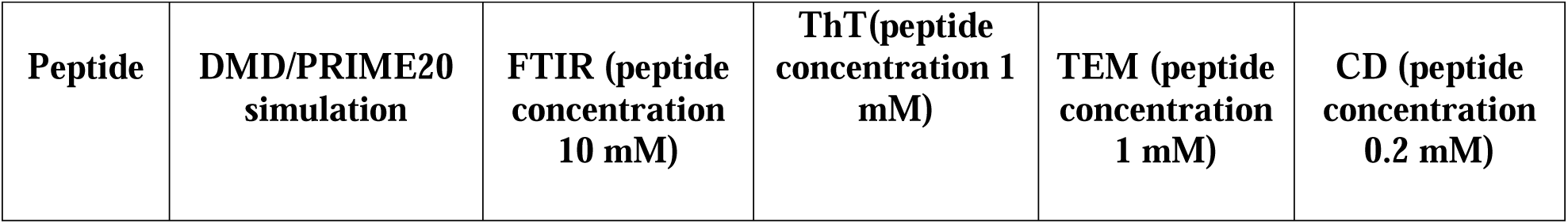
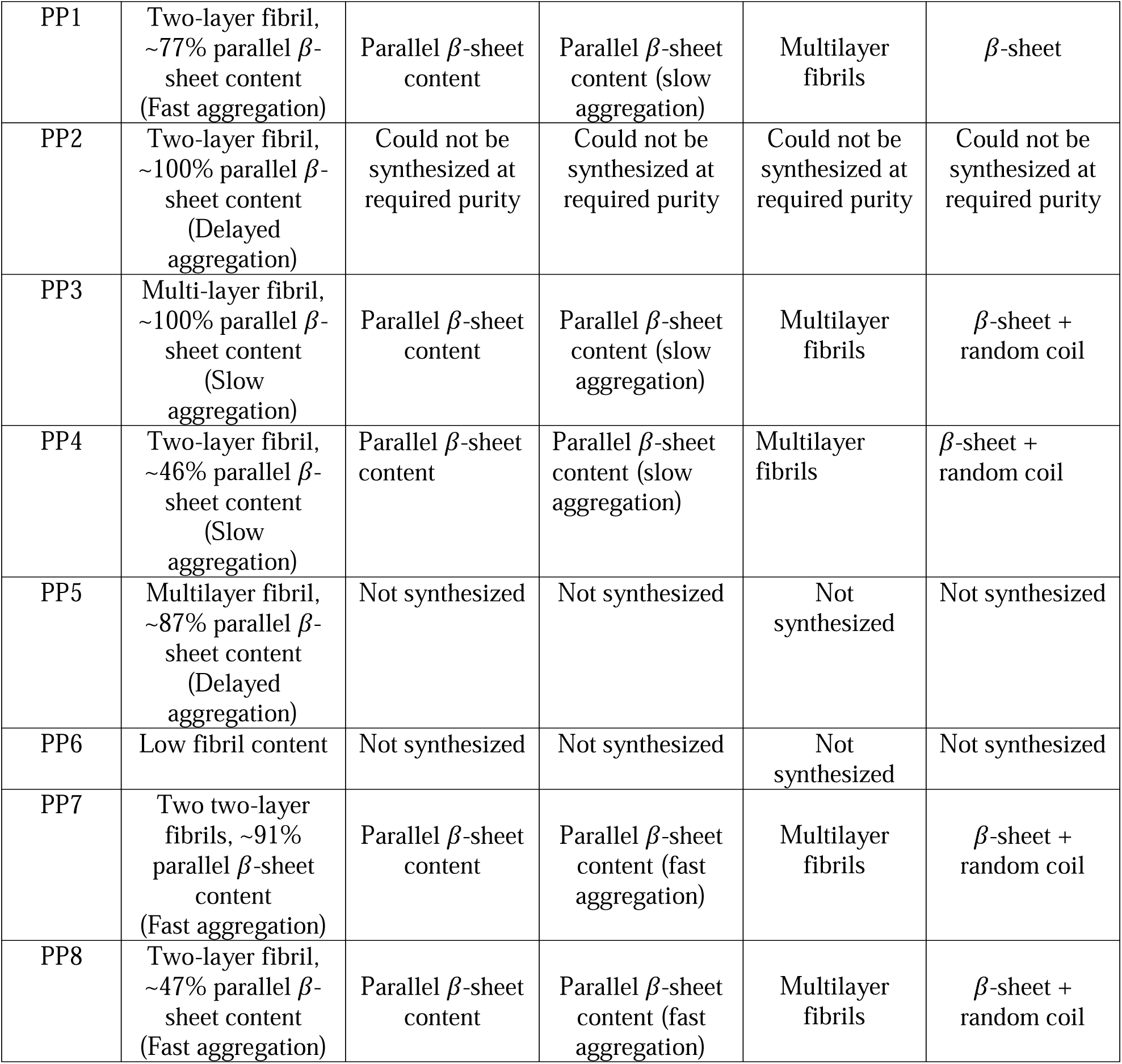
Summary of computational analysis (DMD/PRIME20) and experimental measurements (FTIR, ThT, TEM and CD) of peptides PP1-PP8.

## 3. DISCUSSION AND CONCLUSION

The goal of this work was to identify peptides that self-assemble to form fibrils composed of parallel β-sheets. Inspiration for this work was the Class-1 cross-β spine structure described by Sawaya et al.^24^ that contains two β-sheet layers parallel-oriented β-strands in each layer, and antiparallel-oriented β-sheets between the two layers. Thus far, only one 7-mer peptide, the fibril-forming segment GNNQQNY of the yeast prion protein Sup35, has been identified in the literature as forming amyloid fibrils of the 1^st^ class in experiments. As a first step towards discovering new Class −1 peptides, we set out to design peptides that form fibrils with parallel β-sheets, regardless of the relative orientation of peptides in neighboring layers, by augmenting the workflow involving PepAD, DMD/PRIME20 simulations and experimental characterization. By using the PepAD algorithm coupled with DMD/PRIME20 simulations, we performed two rounds of designs with three different starting backbone scaffolds (**Conf-1**, **Conf-2** and **Conf-3**) to obtain a library of 7-mer amyloid-forming peptides that could potentially assemble into parallel β-sheets. This work complements our previous study where we identified peptides that assemble into anti-parallel β-sheets as are found in the cross-β spine of the 8^th^ class. Discontinuous molecular dynamics simulations with the PRIME20 force field helped us computationally analyze the self-aggregation kinetics of the peptides identified by PepAD. Five out of the eight peptides were synthesized and experimentally tested, and all of them aggregated to form parallel β-sheets. Experimentally, aggregation was detected by observation of nanofibers in TEM images, measurement of CD curves consistent with β-stand secondary structure, positive ThT binding as detected by fluorescence, and FTIR spectra reporting parallel organization of β-sheets.

Although we were successful in showing that computational designs can control organization of β-strands within β-sheets, there are experimental observations that are not readily interpretable based on outputs of PepAD and DMD/PRIME20. First, the ThT measurements indicate a large peptide-to-peptide variation in assembly kinetics, with PP7 and PP8 exhibiting maximal ThT fluorescence in ∼1-2h and PP3 and PP4 showing continuous increase in ThT fluorescence intensity over the course of 72h. Based on DMD/PRIME20 simulations, we had not anticipated large variation in assembly kinetics between different peptides. Second, nanofiber thicknesses varied (Supplementary Figure S4), but all observed thicknesses were far larger than the expected dimension based on 2 β-sheet layers (the interlayer distance between the two sheets from atomistic molecular dynamics simulations is ∼ 4-5Å). Therefore, all the peptides observed to assemble form β-sheet interfaces that were not anticipated in the PepAD algorithm.

To summarize, we now have a computational workflow, PepAD algorithm & DMD/PRIME20 which can output novel sequences that form a desired organization of β-sheets. Thus far we have succeeded in designing peptides that control assembly into parallel or antiparallel β-sheet structures. The predicted structures have been experimentally tested in our previous and current work. For our future work, we could experimentally probe the stacking of β-sheets to validate the predictions of the computational workflow. Additionally, the computational workflow could be developed to discourage the multi-layer fibril formation observed experimentally.

## 4. METHODS

### 4.1 Peptide Assembly Design Algorithm

The *Pep*tide *A*ssembly *D*esign (PepAD) algorithm is a Monte Carlo (MC)-based search procedure to discover peptides that can self-assemble to form supramolecular architectures. In this work, we have focused on designing peptides that self-assemble into the parallel peptides. The procedure is described briefly below.

(1) *<u>Generate input peptide backbone scaffold:</u>* A backbone-scaffold of a reference peptide is required to start the design process. In this work, a peptide scaffold corresponding to the Class 1 cross β-spine (2-layer *β*-sheet structure with 2 parallel-oriented *β*-strands in each layer and antiparallel oriented *β*-strands between the two layers) is generated.
(2) <u>Compute score of initial peptide backbone scaffold:</u> The tendency of the initial peptide backbone scaffold to self-aggregate into a well-organized amyloid-like structure is evaluated using, ***Г**_score_*, a score function that considers the binding affinities between the neighboring chains on the peptide backbone scaffold (Δ*G_binding_*), and the intrinsic aggregation propensities of the individual peptides(*P_aggregation_*). The ***Г**_score_* is defined to be as follows:

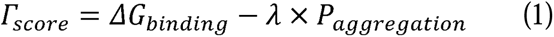 where λ is a weighting factor that adjusts the relative importance of the intrinsic aggregation propensity of the peptides during the sequence evolution.
(3) *<u>Iteration of peptide sequence change moves:</u>* The PepAD algorithm performs 10,000 evolution steps and generates variants of the reference peptide by performing two kinds of trial moves, *viz.* (i) *residue mutation* in which an old residue on all of the peptide chains is randomly chosen and replaced by a new one of the same residue type (hydrophobic, polar, charge, and other); (ii) *residue exchange* in which two residues on all of the peptide chains are randomly chosen and swapped, regardless of their residue type.
(4) <u>*Evaluate score Г_score_ of new peptide sequence:*</u> The ***Г**_score_* for the newly generated peptide sequence draped on the backbone scaffold is evaluated.
(5) *<u>Monte Carlo Metropolis Algorithm:</u>* The Monte Carlo Metropolis algorithm is used to accept or reject new trial peptides.

More details regarding the PepAD algorithm and *Г_score_* can be found in our previous work^27^. The development of the PepAD algorithm has been inspired by our previous work on designing peptides that bind to biomolecular targets using a *Pep*tide *B*inding *D*esign (PepBD) algorithm^35–40^.

### 4.2 Discontinuous Molecular Dynamics Simulation and PRIME20 Model

Discontinuous molecular dynamics (DMD) simulation with the PRIME20 force field have been used to study the fibrilization kinetics of designed peptides by the Hall group. DMD is a fast alternative to traditional molecular dynamics simulations in which the interaction between two particles is modeled with a discontinuous potential, such as hard-sphere, square-well, or square-shoulder potentials. The PRIME20 model is an implicit-solvent coarse-grained protein force field developed in the Hall group that was specifically designed for simulating peptide aggregation with DMD. In the PRIME20 model, each amino acid is represented by three backbone spheres (NH-, C_α_H-, and CO-) and one side chain sphere (R-). Each side chain of the 20 natural amino acids is assigned a unique size, atomic mass, and C_α_-R bond length. Details of the DMD simulations and PRIME20 model are described in our earlier work^28–33^.

In this work, DMD/PRIME20 simulations of the PepAD generated peptides (PP1-PP8) were conducted at T=296K, 303K and 310K for 5 μs for the preliminary screen. The temperature at which a peptide showed highest fibril formation was studied extensively by simulating that peptide for three runs, each at 12 μs. For each *in-silico* peptide system, 48 peptides are placed into a cubic box with edge lengths of 200.0 Å, to achieve a peptide concentration ∼10 mM. In each run, the peptide system starts from a random-coil state. The DMD simulations were carried out in the canonical ensemble. The Andersen thermostat is implemented to maintain the simulation system at the desired temperature. Snapshots of the final simulated structures are obtained using the VMD 1.9.3 software.

### 4.3 Experimental Assessment of Self-Assembly

The sequences output by PepAD were experimentally tested by negative-stain transmission electron microscopy (TEM), Fourier-transform infrared spectroscopy (FTIR), Thioflavin-T Fluorescence and Circular Dichroism (CD) using the methods we detailed previously^23^. All peptides were imaged using negative-stain transmission electron microscopy at a peptide concentration of 1mg/ml in deionized water (DI water). FTIR measurements were conducted after a minimum of 72h post assembly at a concentration of 10mM in DI water. Additionally, we performed Circular Dichroism experiments at a concentration of 0.2mM in DI water. Thioflavin-T Fluorescence experiments were at a peptide concentration of 2mM over an assembly period of 72h.

## ACKNOWLEDGMENTS

The authors in this work would like to thank the National Science Foundation (Award No. OAC-1931430) for financial support. This work used the Extreme Science and Engineering Discovery Environment (XSEDE), supported by National Science Foundation grant number ACI-1548562. SS and CKH thank San Diego Supercomputer Center (SDSC) for providing computing time.

## DATA AVAILABILITY

Data is available in the manuscript and/or supporting information. The PDB files of starting conformations (Conf-1, Conf-2 and Conf-3) and PepAD output files of PP1-PP8, PDB output files and beta-sheet content from DMD/PRIME20 simulations of PP1-PP8 and analyis codes are available at:
https://github.com/CarolHall-NCSU-CBE/Parallel-self-assembling-peptides.

## Supporting Information for Publication

### Supplementary Figures

**Figure S1.**
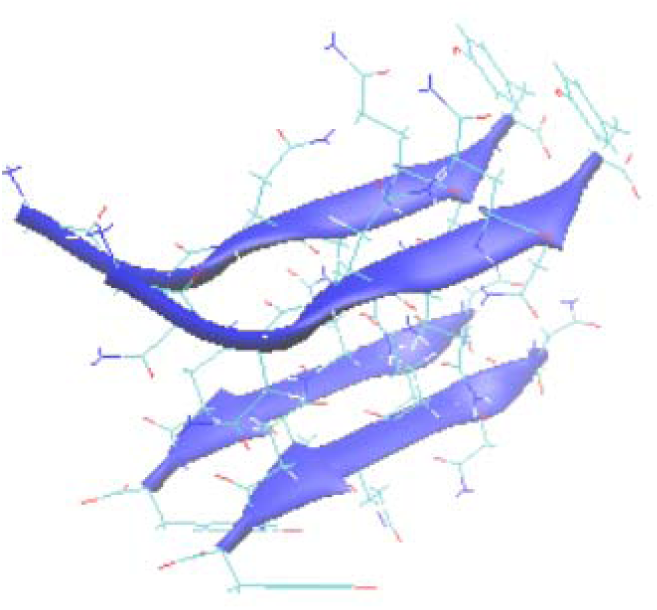
Class 1 cross-*β* spine containing 4 GNNQQNY peptides.

**Figure S2.**
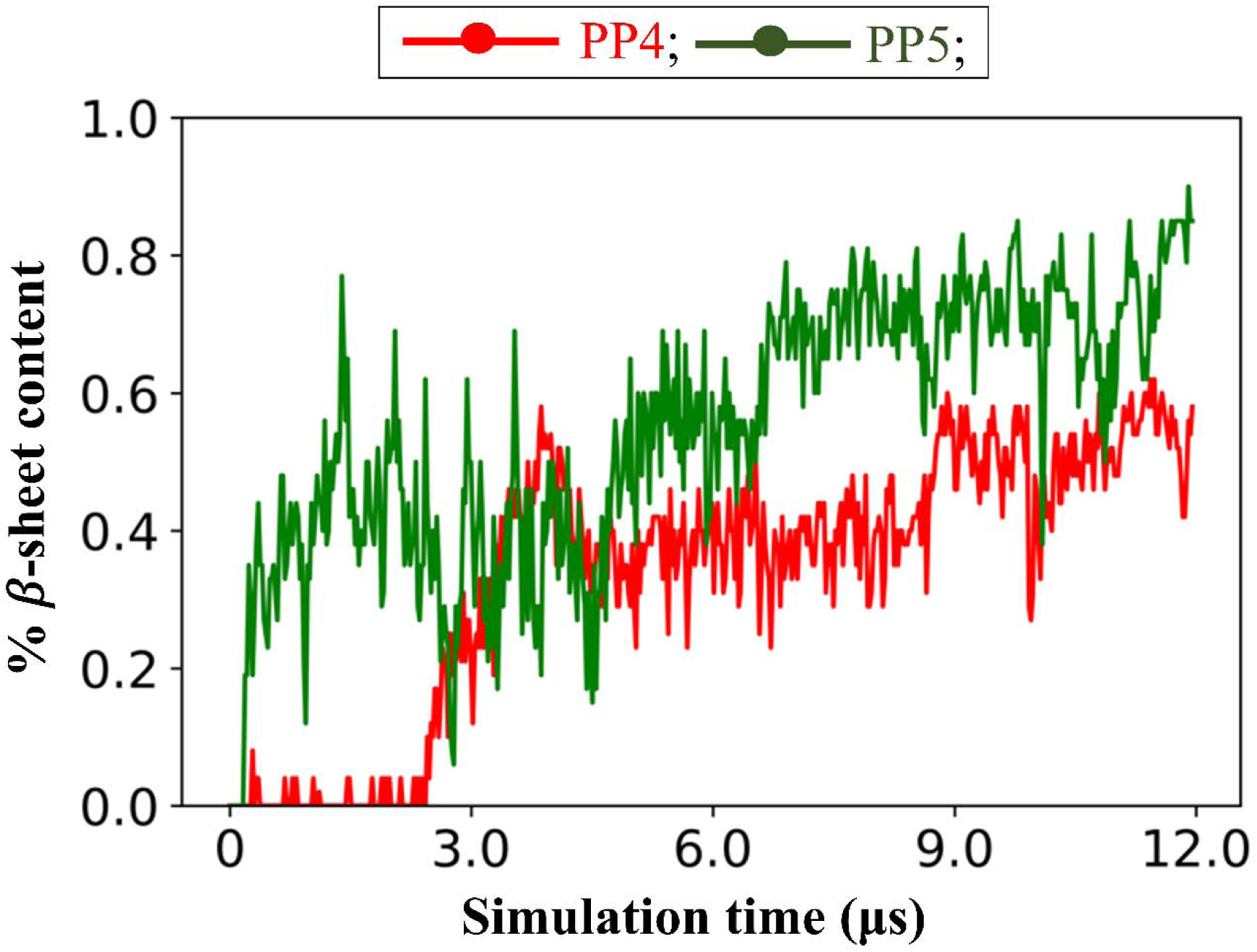
Plot of *β*-sheet content v/s simulation time describes the self-aggregation kinetics of peptide PP4: GAIDWVK and PP5: ADKVMFV.

**Figure S3.**
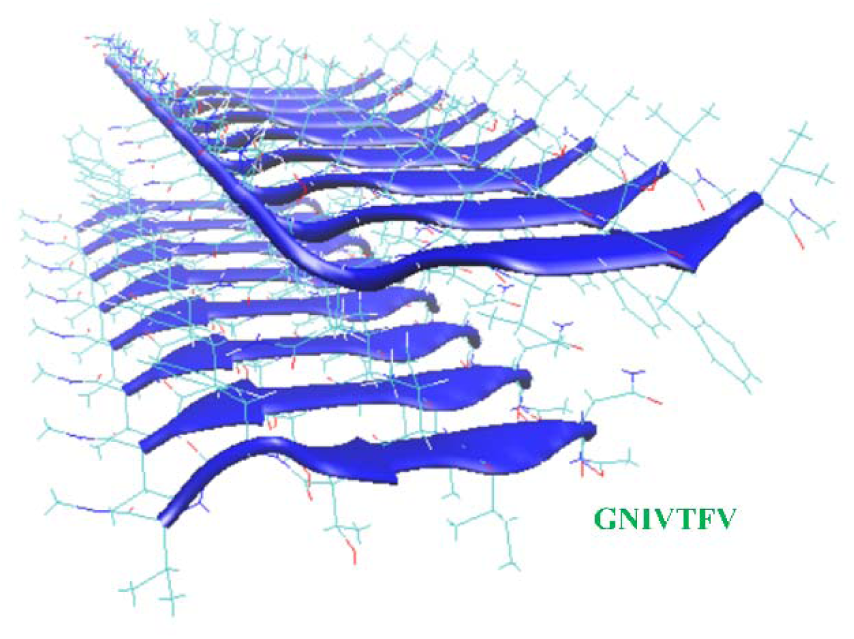
Class 1 cross-*β* spine structure containing 16 GNIVTFV (PP2) peptides.

**Figure S4.**
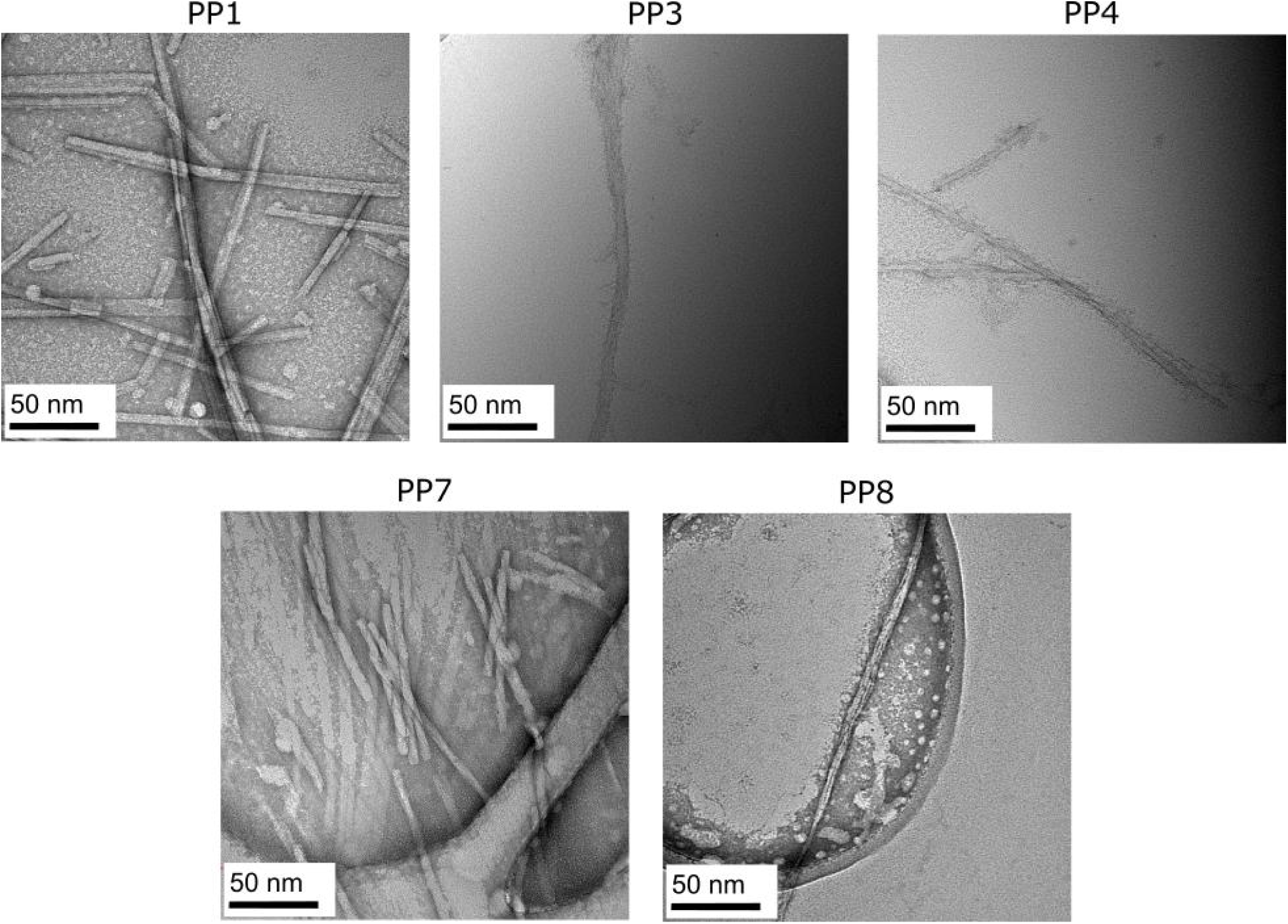
Transmission electron micrographs of negatively stained samples prepared one day following peptide dissolution at 1 mg/ml.

**Figure S5.**
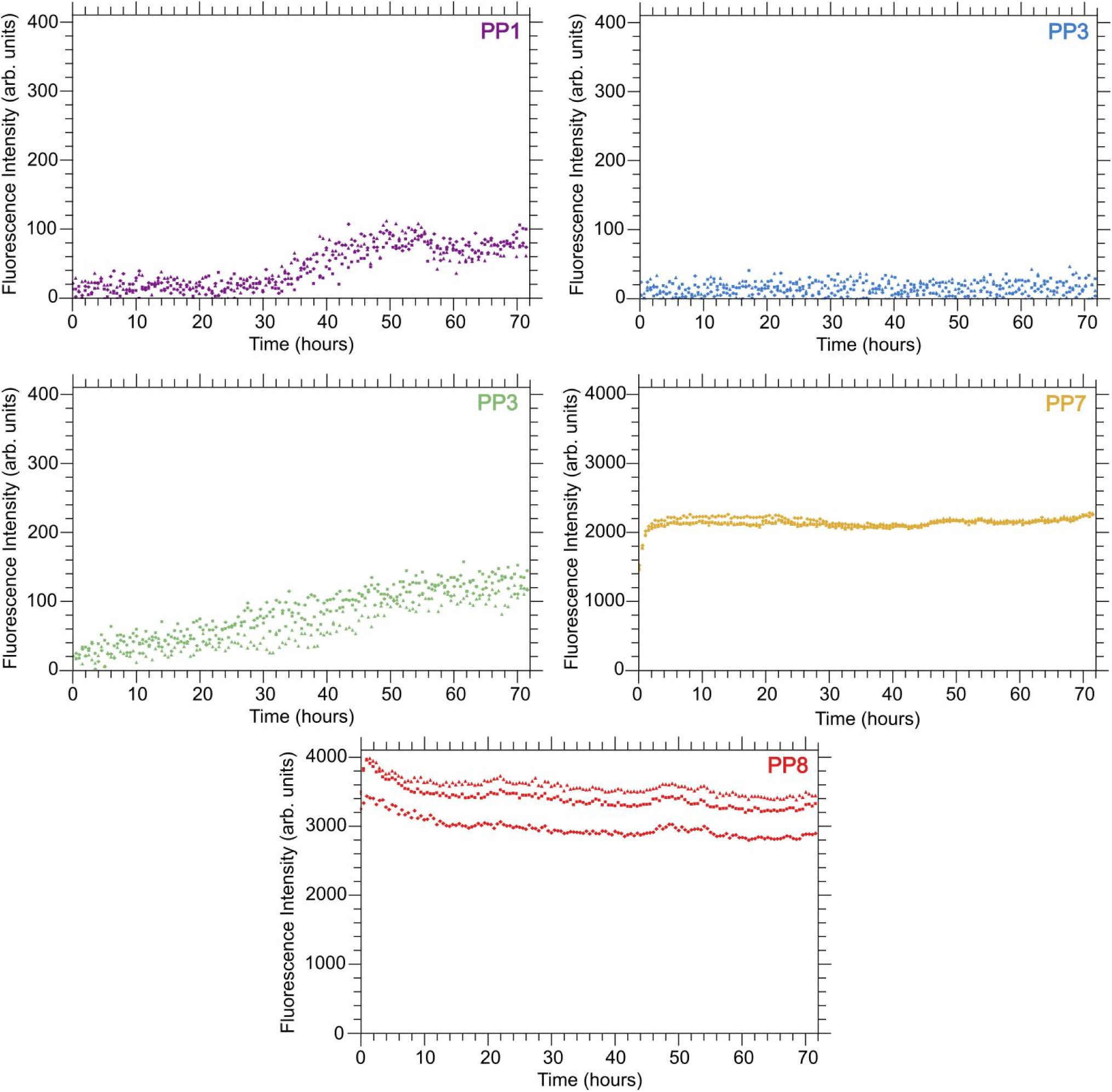
ThT fluorescence replicate curves for each of the five peptides predicted to aggregate.

### Peptide Synthesis

Peptides (PP1, PP2, PP3, PP4, PP7 and PP8) were purchased from CPC Scientific, Inc. (Sunnyvale, CA). The peptides received at > 95% purity were used in experiments without modification.

### Thioflavin-T fluorescence

Thioflavin-T (ThT) measurements were conducted using the procedure from our previous work^1^. Peptides were dissolved to a concentration of 2mM total peptide, 0.08mg/mL ThT, and DI water before being added to a black 96-well plate (Thermo Scientific Nunc). Peptide samples were analyzed using a BioTek Synergy H4 Microplate Reader (excitation 450nm, emission 482nm, slit bandwidth 9nm), and fluorescence intensity was recorded over 72h. We performed ThT fluorescence measurements in triplicate, with the means of the samples reported in the main text.

### Fourier Transform Infrared Spectroscopy

Fourier transform infrared spectroscopy (FTIR) was performed using protocol developed in our previous study^1^. These measurements were conducted on Thermo Scientific Nicolet 6700 spectrometer with an attenuated total reflection (ATR) accessory. Peptide solutions prepared at 10mM were spotted onto the ATR accessory, and an average over 64 scans was collected after an assembly period of 72h. As PP3 and PP4 assembled slower as compared to rest of peptides, FTIR scans were collected after 6 days of assembly.

### Transmission Electron Microscopy

Peptide solutions with DI water at 1mg/ml were prepared and assembled for minimum of 24h before measurements. Transmission Electron Microscopy (TEM) was conducted with a protocol similar to our previous work^1^. As PP3 and PP4 assembled slowly, an assembly period of 6 days was employed prior to TEM measurements.

### Circular Dichroism Spectroscopy

Peptides were dissolved in water at a concentration of 0.2mM in DI water (De-ionized). Subsequent to dissolution, concentration was verified using UV/Vis absorbance. Following a 2h assembly period, CD was measured at room temperature with a Chirascan^TM^-plus spectrometer (Applied Photophysics, Ltd.), following baseline correction with DI water without peptide. Quartz cuvettes with a 0.1mm path length were used. The protocol established for our previous study was employed to conduct measurements.

### Comparison of secondary structure predicted by BeStSel and our current study

BeStSel is a web server-based platform for the determination of secondary structure from circular dichroism spectra developed by Micsonai et al.^2^ All tested peptides exhibit parallel *β*-sheet structures as evidenced by biophysical characterization techniques like negative-stain transmission electron microscopy (TEM), Fourier-transform infrared spectroscopy (FTIR), Thioflavin-T Fluorescence and Circular Dichroism (CD). We compared our interpretation of Circular Dichrosim experiments to the secondary structure predicted by BeStSel when data from the experiment was input into it. We suggest that results are unreliable when the peptides tested have heterogenous structure.

**Table S1.** Summary of secondary structures predicted by BeStSel and secondary structures determined by our characterization techniques.

| Peptide | BeStSel | Current Study: CD interpretation | Result from current study: All techniques |
| --- | --- | --- | --- |
| PP1 | 100% parallel β-sheet | β-sheet | Parallel β-sheet |
| PP3 | 12.2% helix, 32.1% antiparallel β-sheet, 5.7% parallel β-sheet, 49.9% other | β-sheet + random coil | Parallel β-sheet (slow aggregation) |
| PP4 | 20.1% helix, 64.5% antiparallel β-sheet, 15.3% other | β-sheet + random coil | Parallel β-sheet (slow aggregation) |
| PP7 | 44.3% helix, 55.7% antiparallel β-sheet | β-sheet + random coil | Parallel β-sheet |
| PP8 | 9.1% helix, 9.1% antiparallel β-sheet, 18.3% parallel β-sheet, 57.2% other, 6.4% turn | β-sheet + random coil | Parallel β-sheet |

## Notes

https://github.com/CarolHall-NCSU-CBE/Parallel-self-assembling-peptides

### Competing Interest Statement

The authors have declared no competing interest.

